# Slow wave sleep supports the reorganisation of episodic memory networks

**DOI:** 10.1101/2025.03.24.644966

**Authors:** Simon Faghel-Soubeyrand, Polina Perzich, Bernhard P. Staresina

**Affiliations:** Department of Experimental Psychology and Oxford Centre for Human Brain Activity, Wellcome Centre for Integrative Neuroimaging, Department of Psychiatry, University of Oxford, Oxford, United-Kingdom

**Keywords:** episodic memory, sleep, memory consolidation, memory transformation, alpha rhythms, source analysis

## Abstract

Models of memory consolidation propose that newly acquired memory traces undergo reorganisation during sleep. To test this idea, we recorded high-density electroencephalography (EEG) during an evening session of word-image learning followed by immediate (pre-sleep) and delayed (post-sleep) recall. Polysomnography was employed throughout the intervening night, capturing time spent in different sleep stages. Using source-reconstructed time-frequency analysis, we first replicated the effect of alpha power decreases for successful relative to unsuccessful recall, emerging between 700 and 1500 ms after cue onset and spanning medial and lateral temporal lobe regions as well as posterior parietal cortex. Directly contrasting successful post-sleep vs. pre-sleep recall revealed a shift of alpha power decrease from parietal towards anterior temporal lobe (ATL) after sleep. Critically, time spent in slow wave sleep (SWS) during the intervening night not only predicted the extent of memory retention, but also correlated with the shift to ATL recall effects. Finally, brain-wide functional connectivity profiles during successful recall pointed to a marked overnight reorganisation of memory networks, with the extent of reorganisation again predicted by time spent in SWS. Together, these findings link SWS to the consolidation and functional reorganisation of episodic memory networks.

## Introduction

Our memories of past events and experiences are not static, but change in both strength and quality as a function of time (Dudai et al., 2015; Gilboa & Moscovitch, 2021). Models of systems consolidation propose that these phenomenological changes are accompanied by a gradual shift from hippocampal to neocortical involvement during recall (Born & Wilhelm, 2012; Frankland & Bontempi, 2005; Klinzing et al., 2019; McClelland et al., 1995; Rasch & Born, 2013). By strengthening cortico-cortical connectivity, this transfer is thought to enhance memory stability and promote the integration of new information into existing knowledge networks.

Crucially, sleep constitutes an optimal state for systems consolidation to occur, as the brain is sheltered from external stimulation during this period (Stickgold & Walker, 2005; Winocur & Moscovitch, 2011). Indeed, a large body of animal and human work has linked non-rapid-eye-movement (NREM) sleep and concomitant sleep rhythms (slow oscillations, spindles and ripples) to the purported hippocampal-neocortical dialogue supporting memory consolidation (Buzsáki, 1998; Diekelmann & Born, 2010; Frankland & Bontempi, 2005; Staresina, 2024). However, the influence of consolidation on memory recall dynamics, along with the specific role of sleep in this process, remains poorly understood. Prior fMRI work has revealed a shift in recall networks from hippocampus (immediate recall) to medial prefrontal cortex (delayed recall after 24h) (Takashima et al., 2009). In parallel, functional connectivity increased between fusiform cortex and other cortical regions, but decreased between fusiform cortex and hippocampus, pointing to a shift towards cortico-cortical connectivity for recall of consolidated memories. Nevertheless, whether this shift in recall networks was directly linked to overnight sleep remained unclear. Another study used Magnetoencephalography (MEG) to examine differences in recall dynamics for recent (1h) and remote (25h) memories and found strong connectivity changes between learning-related cortical regions and the anterior temporal lobe (ATL; Nieuwenhuis et al., 2012). While these findings ascribe a special role to the ATL and its proposed role as a semantic hub (Lambon Ralph et al., 2010; Patterson et al., 2007; Ralph et al., 2017) in recall of consolidated memories, there was again no direct link between these network changes and overnight sleep characteristics.

With regard to recall dynamics, recent work has highlighted alpha rhythms (8–12 Hz) as a viable indicator of recall success (Griffiths et al., 2019; Martín-Buro et al., 2020). Specifically, decreases in alpha power have been consistently linked to successful memory recall, localised to the canonical retrieval network that includes medial temporal, posterior parietal and lateral prefrontal regions (Griffiths et al., 2019; Hanslmayr et al., 2012; Martín-Buro et al., 2020; Treder et al., 2021). Notably, shifts in the spatial distribution of recall-related alpha decreases over time may provide insights into how memories are reorganised within cortical networks.

Here, we assess how sleep may support the reorganisation of memory recall networks. We recorded high-density EEG during a word-image learning task followed by two recall phases, one pre-sleep and one post-sleep, with intervening sleep monitored via Polysomnography. Using source reconstruction of brain signals and connectivity analyses during the recall phase, we specifically investigate shifts in the spatial distribution of memory networks following sleep. We also examine whether these neural shifts relate directly to time spent in different sleep stages.

## Methods

### Participants

A total of 24 participants (5 male, M_age_ = 22.83) were recruited for this study and compensated financially or with course credits. All participants had normal or corrected-to-normal vision. Participants were screened for sleep-related issues using the Pittsburgh Sleep Quality Index (PSQI, Buysse, Reynolds, Monk, Berman, and Kupfer, 1989) and excluded if they had a score of 5 or above on the PSQI, if they were taking any sleep modifying medication, did night shifts at present/in the past year, if they were smokers, or if they had a history of neurological, psychiatric, or sleep disorders. The research was approved by the University of Oxford’s ethics committee (approval code: R75681/RE002). All participants provided written informed consent prior to participation.

### Stimuli and Task

Participants completed two separate sessions of a word-image memory task, with a minimum interval of 7 days (n=21) and a maximum interval of 8 days (n=3) between them. In ‘object sessions’, the images associated with words were either a car or a guitar. In ‘scene sessions’, images were either a house or a corridor. The order of object and scene sessions was counterbalanced across participants. Within a session, the two object or scene images were paired with 100 trial-unique verbs. Images were sampled from the fLoc functional localiser package (Stigliani et al., 2015). MATLAB and Psychtoolbox (Brainard, 1997) were used for the experimental code and stimulus preparation. Stimuli were displayed on a 2500 Intel UHD Graphics 770 monitor with a resolution of 1920 × 1080 pixels and a refresh rate of 60 Hz. Participants were seated 60 cm away from the monitor.

The task consisted of an encoding (learning) phase followed by two retrieval phases, one before sleep and one after sleep.

### Encoding

During the encoding phase, participants were shown a single word alongside an image of an object or scene and were asked to evaluate the plausibility of the association in a 2AFC procedure. For example, the word “drive” paired with an image of a car would likely be rated plausible, whereas the same word paired with an image of a guitar would likely be rated implausible. Note however that plausibility ratings were subjective. Each trial began with a fixation cross presented for 1250 ms, with a variable jitter of ±250 ms. The word-image pair then appeared on the screen for a minimum of 2.5 seconds, with a maximum response window of 10 seconds. After 2.5 seconds, a response cue remained on the screen until the participant responded or until the 10-second response window elapsed. Participants completed 100 encoding trials, with a self-paced break after every five minutes of encoding.

### Retrieval

During the retrieval phase, participants were shown the previously presented words and had to indicate which image had been associated with the word during encoding (cued recall). Each retrieval trial was presented in the following sequence: a fixation cross was displayed for 1250 ms, with a ±250 ms jitter, followed by the word cue presented for a minimum of 2.5 seconds and up to a maximum of 5 seconds. Participants could indicate their response as soon as the response display appeared (after 2.5 seconds). The response window remained open until the participant responded or until the maximum duration of 5 seconds was reached. 50 encoded word-image associations were randomly chosen for pre-sleep retrieval, with the remaining 50 for post-sleep retrieval. Note that participants also had a ‘don’t know’ response option and were encouraged to use that option instead of guessing the target image. For EEG analyses, incorrect and ‘don’t know’ responses were pooled into a single ‘unsuccessful recall’ condition.

After each night of sleep, participants were woken up by the experimenter at approximately 7:30 AM and were given a few minutes to fully awaken. The impedances of the EEG electrodes were then checked, with adjustments made for any electrodes exceeding 20 kΩ. This was followed by an attention task designed to reduce sleep inertia, as described below.

### Attention task

A psychomotor vigilance task (PVT) was included as an offline period prior to the two retrieval phases (one in the evening and one in the morning). The PVT lasted three minutes per session and required participants to monitor a white fixation cross on a black screen. At unpredictable intervals, a numerical counter appeared at the centre of the screen, which increased in milliseconds until the participant responded by pressing a designated key. Reaction times were recorded, with faster responses indicating higher alertness and slower responses reflecting lapses in attention. If no response was given within two seconds, a message was displayed prompting participants to remain attentive. Feedback on reaction times was provided following each response. The task included random inter-stimulus intervals ranging from 10 to 15 seconds.

### Behavioural analysis and dimensionality reduction

We calculated an adjusted recall accuracy score by subtracting the number of incorrect responses from the correct responses and then dividing the result by the total number of correct, incorrect, and ‘don’t know’ trials. To assess memory retention, we measured the change in adjusted recall accuracy from pre-sleep to post-sleep (i.e., subtracting pre-sleep from post-sleep), reflecting the extent to which memory was retained overnight. A similar score was derived from recall response time by calculating the mean response time for correct trials and assessing the difference between post-sleep and pre-sleep response times. We then applied Principal Component Analysis (PCA) to the distributions of adjusted memory accuracy and response time change, extracting a single memory retention score based on the first component, which accounted for the highest variance between the two performance measures. This memory retention score was used for all analyses relating memory retention to i) sleep and ii) brain imaging data.

### EEG acquisition

Scalp electroencephalography (EEG) was recorded using a Brain Products 64-channel EEG (10-20 system) system with a sampling rate of 500 Hz. 6 channels were re-purposed for Polysomnography – 2 for mastoid recordings, 2 to measure electromyography (EMG), and 2 to record eye movements (electrooculography, EOG). Thus, a total of 58 channels were left for recording of scalp EEG.

### Sleep scoring

Sleep stages were first scored based on established detection algorithms (Guillot & Thorey, 2021; Vallat & Walker, 2021), using channels C3-M2, E1-M2, and EMG. Whenever these two sleep staging algorithms were not in agreement, manually scoring was employed according to the AASM guidelines (Iber et al., 2007). The average of sleep duration/proportion across the 2 sessions per participant was used for further analyses.

### EEG preprocessing

The Fieldtrip toolbox (Oostenveld, Fries, Maris, & Schoffelen, 2011) was used for data preprocessing and analysis. We first rejected eye blink artifacts from continuous EEG data using Independent Component Analysis (ICA). For each participant and session, we manually inspected the topography and time course of each component to identify and reject those associated with artifacts. The remaining components were then back-projected to reconstruct the cleaned EEG signal. This was followed by bad-channel identification and interpolation using the average of neighbouring electrodes. Raw EEG recordings were then filtered using a 0.5 Hz high-pass filter to remove low-frequency drifts. Lastly, data were re-referenced to the common average of all channels.

### Source localisation

Source localisation was performed in a similar way as in Treder et al. (2021). Electrode positions were defined using our 64-channel EEG cap and manually realigned to a standard Fieldtrip head model. The head model was constructed using a boundary element model (BEM), and the standard MNI brain template was used for anatomical reference. A spatial filter was estimated for each grid point in the source model using either linearly constrained minimum-variance beamformers (LCMV) for Time Frequency Representation analyses (with regularisation λ = 10%) or partial canonical correlation (PCC) for connectivity analyses (again with regularisation λ = 10%). Source activity was projected onto a 10-mm grid defined in MNI space with 3,294 grid points inside the brain. For region-of-interest (ROI) analyses in source space, we used anatomical masks provided by the Automated Anatomical Labeling (AAL) atlas (Tzourio-Mazoyer et al., 2002)

### Time frequency analysis

Time-frequency representations of EEG data were computed using a sliding window Fourier Transform approach. The window length was set to five cycles of a given frequency and the windowed data segments were multiplied with a Hanning taper. The time window for computation of TFR ranged from-3.5 to 5.5 seconds relative to the event onset (i.e., the cue) with power computed in steps of 50 ms. The power spectra of single-trial data were extracted from-.5 s to 2.5 s relative to cue onset, and examined for outliers based on the distribution of spectral power across trials (using MATLAB’s *isoutlier* function). Outlier values were excluded from further analysis. Data were then normali*s*ed using a z-scoring approach across time and trials dimensions. Then, condition-specific TFRs were computed by averaging trial-level power spectra. Statistical analyses were then performed on participant-and condition-wise TFRs.

### Connectivity analysis

Connectivity analysis was performed to investigate interactions between brain regions during the retrieval phase. Source-level data were estimated using beamforming with partial canonical correlation (PCC), projecting sensor-space Fourier coefficients onto a predefined source model. Individual trials were retained to enable connectivity estimation. Anatomically defined regions of interest (ROIs) were identified using an MNI-based AAL atlas, and for each ROI, spectral power and momenta were extracted across trials. Connectivity measures were computed between these 24 ROIs using imaginary coherence, which mitigates the effects of volume conduction by considering only non-zero phase-lagged interactions (Nolte et al., 2004). Trial counts were equated between conditions through random subsampling, and outliers were excluded based on power distributions as described above. Each participant’s connectivity values were computed across 20 resampling iterations, and averaged for the final ‘connectomes’. Participant-and condition-specific connectivity matrices were statistically compared at the group level. To compute the ‘connectome distance’ (i.e., changes in brain-wide connectivity-profiles) between pre-sleep and post-sleep recall, we used 1 – the cosine similarity between the two 24 × 24 connectivity matrices for each participant. This connectome distance was then correlated with measures of interest, i.e., sleep stage duration/proportion and memory retention.

### Statistical tests

For TFR analysis in source space, dependent-samples *t*-test were used to compare spatiotemporal maps of alpha band activity across conditions. The statistical analysis was conducted using a Monte Carlo method with 1000 random permutations to estimate the null distribution of the test statistic, and a cluster-based correction was used to control for multiple comparisons across spatial sources (Maris & Oostenveld, 2007). Clusters were identified based on a cluster-forming threshold of *p* < 0.05 (two-tailed). The results were visualised by interpolating significant clusters onto a structural MRI template, and statistical values were projected onto cortical surface models for topographical visualisation. The statistics for the Time x Frequency x channel analysis (Supplementary Material) were conducted in a similar manner, using dependent-samples *t*-tests and a cluster-based correction method employed to identify significant clusters in time, frequency, and electrode space. First, individual time-frequency points were thresholded at *p* < 0.05 (two-tailed), and neighbouring points were grouped into clusters based on their spatial, temporal, and spectral adjacency. For all correlations with sleep stages and memory, Spearman correlations were used throughout. For correlations with sleep stages, we rejected outliers prior to computing the Spearman-*r*. This yielded three outliers when looking at N3 proportion, and two outliers when looking at N3 duration.

## Results

### Behavioural results

Participants performed two experimental sessions separated by 1 week, with image category (objects, scenes) alternating between sessions (counterbalanced across participants) (see Methods and **Figure 1**). Pooled across both sessions and recall phases, participants gave an average of 63.65% correct target image responses, 8.83% incorrect target image responses, 27% ‘don’t know’ responses, and 0.52% invalid responses (i.e., no button press within the response window). To quantify recall success, we used an adjusted recall score, subtracting the proportion of incorrect target image responses from the proportion of correct target image responses.

**Figure 1.**
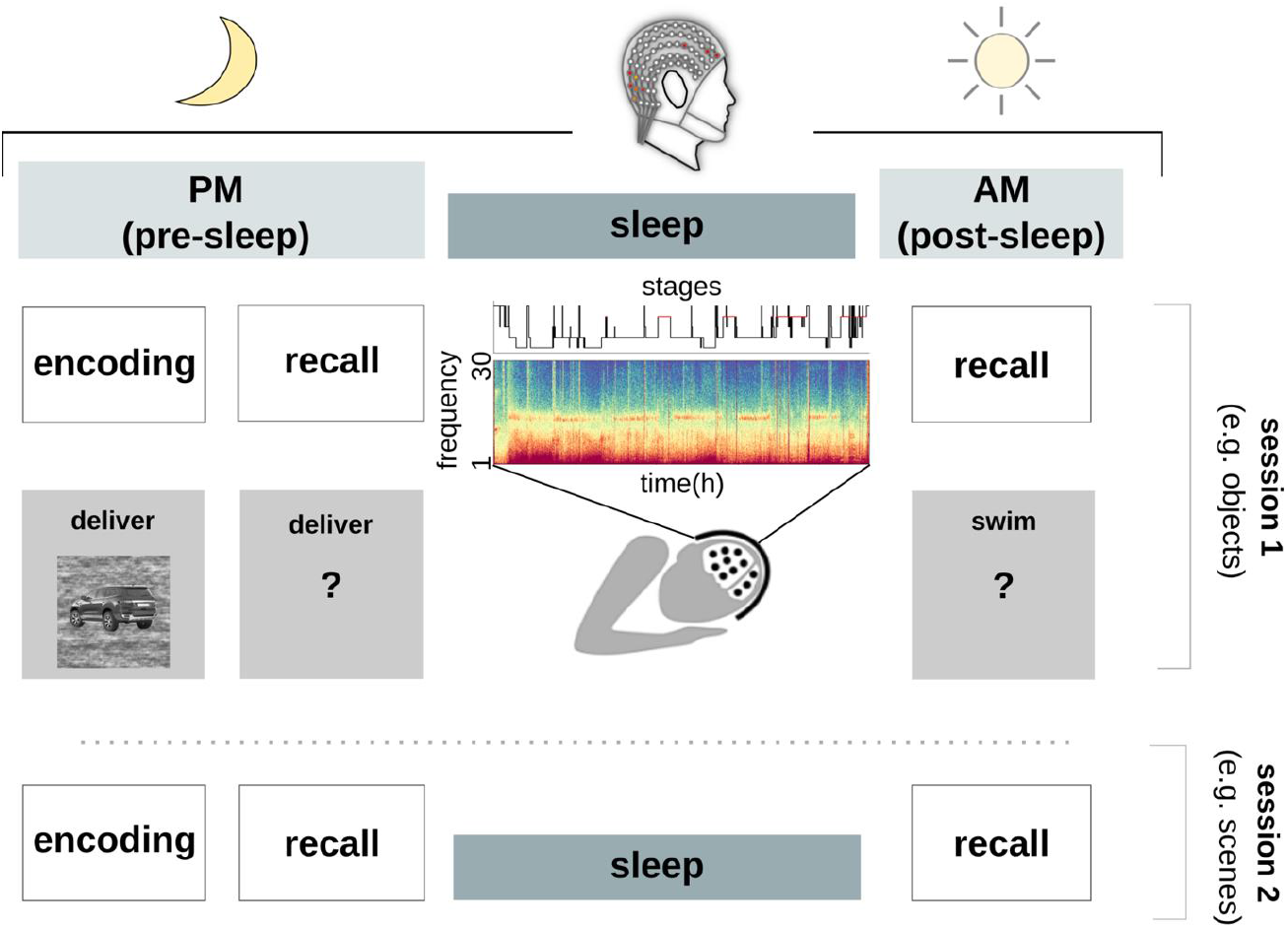
Experimental procedure. Participants completed two sessions of an episodic memory task (at least 1 week apart) including high-density EEG recordings. Each session contained three phases: (i) encoding (learning), (ii) pre-sleep recall (PM), and (iii) post-sleep recall (AM). Throughout the night, sleep was recorded using high-density EEG/PSG. Each session included stimuli from either object (car or guitar) or scene (house or corridor) categories, counterbalanced across participants. Each trial of the encoding task consisted of a unique word-image association, from which a random subset of 50% were tested during pre-sleep recall, with the remaining 50% tested during post-sleep recall.

Pooled across object and scene sessions, participants’ adjusted recall score of dropped from 65.03% (SEM = 4.25) before sleep to 45.21% (SEM = 4.44) after sleep. To statistically quantify the effect of delay and possible effects of image category, we performed a repeated-measures ANOVA on adjusted recall scores including the factors Delay (pre-sleep, post-sleep) and Category (objects, scenes). Results revealed a significant main effect of Delay (F (95) = 18.84, *p* <.0001) and no effect of Category (F (95) = 0.06, *p* =.801), nor a Delay x Category interaction (F (95) = 0.01, p =.944). The effect of Delay on recall scores was paralleled by effects on response times (RTs) for correct recall responses: The mean RT was 3.18 s (SEM = 0.05) before sleep and 3.34 s (SEM = 0.05) after sleep. A repeated-measures ANOVA on RTs during successful recall including the factors Delay (pre-sleep, post-sleep) and Category (objects, scenes) showed a significant main effect of Delay (F (95) = 7.44, *p* =.008), and again and no effect of Category (F(95) = 0.26, *p* =.609), nor a Delay x Category interaction (F(95) = 0.05, p =.818, see also **Figure 3a**). Since Category-type (objects vs. scenes) had no effects on any of the behavioural outcomes, we averaged EEG results across the two experimental sessions per participant for the remainder of the analyses.

### Local and network-level brain dynamics of successful memory recall

We first aimed to investigate the spatiotemporal oscillatory dynamics underlying successful memory recall, collapsing data across pre-and post-sleep recall. At the outset, we calculated the contrast of successful vs. unsuccessful recall trials for all channel x time x frequency combinations. As shown in **Supplemental Figure 1**, we replicated previous findings of decreased alpha power (peaking at 10 Hz) during successful recall, emerging at ∼700 ms post cue onset and showing a topographical peak over left parietal channels (Martín-Buro et al., 2020; Treder et al., 2021). Based on this result, we focused all subsequent analyses on the alpha band (8-12 Hz). Next, we projected the raw EEG signals into source space (see *Source Localisation* in Methods) and computed alpha-band time-frequency representations (TFRs) for successful and unsuccessful recall trials. We then compared the resulting average alpha-band source-level representations in 100 ms time windows (0–2500 ms post-cue onset) using paired t-tests and cluster-based correction for multiple comparison across source voxels (Oostenveld, Fries, Maris, & Schoffelen, 2011). This analysis revealed a set of spatial clusters showing significant alpha power decreases for successful recall emerging from approximately 700 ms and persisting until around 1500 ms after cue onset. The temporal evolution of significant clusters is shown in **Figure 2a**. It followed a progression from medial temporal and parietal regions to engagement of more anterior temporal regions as memory recall unfolds. This dynamic pattern aligns with previous findings that associate alpha decrease with memory access and the reactivation of stored representations (Griffiths et al., 2019; Martín-Buro et al., 2020; Staresina & Wimber, 2019; Treder et al., 2021).

**Figure 2.**
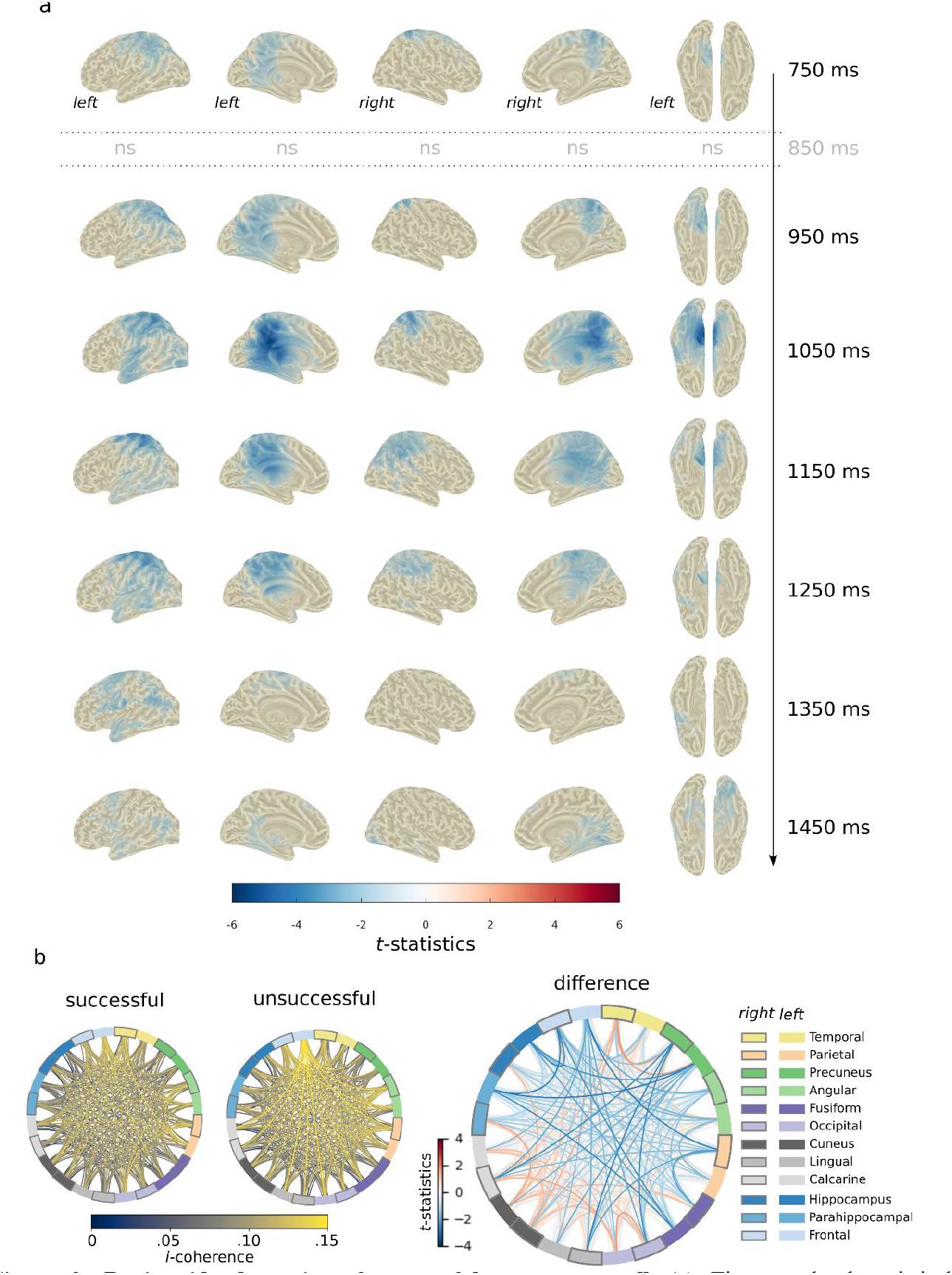
Brain-wide dynamics of successful memory recall. (a) Time-resolved statistical comparison of alpha-band decreases during recall. Cortical surface maps display significant *t*-statistics (cluster-test, *p* <.05, two-sided) contrasting successful versus unsuccessful trials at successive 100 ms time windows, which were significant from 700 ms to 1500 ms post-cue onset. Blue regions indicate significantly stronger alpha decrease for successful trials. (b) Alpha-band functional connectivity networks for successful (left) and unsuccessful (middle) recall, computed using the imaginary part of coherence. The rightmost panel shows the difference(paired *t*-tests) in connectivity between conditions, with blue connections indicating stronger decreases during successful trials and red connections indicating stronger decreases during unsuccessful trials.

We next examined network connectivity associated with successful recall. Following our previous observation of memory-related alpha power decreases we computed alpha-band connectivity for successful and unsuccessful recall within a one-second time window of interest (starting at the first point of significant alpha power differences, i.e., at 700ms, again first collapsing pre-and post-sleep recall). The extension to a 1 s window yielded a frequency resolution of 1 Hz. The imaginary part of coherence is a robust connectivity measure that focuses on phase-lagged interactions, which minimizes the influence of volume conduction and highlights genuine functional connectivity (Nolte et al., 2004). To enhance the interpretability of the resulting memory network, the analysis was restricted to cortical and subcortical regions of interest derived from the AAL atlas (Tzourio-Mazoyer et al., 2002). The corresponding connectivity plots are shown in **Figure 2b**. Similar to Figure 2a, which identified local alpha power reductions, the difference plot in Figure 2b contrasts successful vs. unsuccessful recall trials, now highlighting differences in brain-wide alpha-band connectivity between the two conditions. This analysis revealed a wide-ranging modulation of functional connectivity linked to successful memory recall. Rather than interpreting individual, pairwise connectivity patterns, we consider the entire connectivity structure (‘connectome’) when querying overnight changes in recall network profiles in subsequent analyses. Together, these results show how successful recall is reflected in both localised and network-level alpha decreases.

### Deep sleep predicts memory retention

Having identified local and global patterns of alpha power linked to recall performance, we next strived to elucidate the effect of overnight sleep on these patterns. To this end, we first obtained a single behavioural index of overnight memory retention per participant. That is, given that recall accuracy and response times (RT) were similarly affected by immediate vs. delayed recall, we used principal component analysis on both metrics, revealing that the first component explained 89.77% of the variance shared between accuracy and RT changes across the night. We henceforth use this component score as our behavioural measure of memory retention.

Absolute time and proportions spent in different sleep stages are shown in Table 1. Importantly, correlations between sleep stages and memory retention revealed one significant relationship: a positive correlation between N3 proportion and memory retention (*r* = 0.65, *p* = 0.001). This correlation also held when using the absolute duration of N3 sleep (*r* =.52, p =.013). No significant correlation was found between memory retention and any other sleep stage (all *ps* >.1).

**Table 1.** Summary of sleep stage duration and proportion.

|  | mean | std-dev |
| --- | --- | --- |
| N1 (minutes) | 31.42 | 13.7 |
| N2 (minutes) | 254.92 | 29.35 |
| N3 (minutes) | 82.79 | 20.0 |
| REM (minutes) | 100.81 | 18.94 |
| N1 (%) | 6.74 | 3.06 |
| N2 (%) | 54.24 | 4.41 |
| N3 (%) | 17.67 | 4.34 |
| REM (%) | 21.36 | 3.13 |

### Overnight sleep reorganises memory networks

We showed that successful recall is associated with alpha-band decreases occurring approximately.7 to 1.5 seconds after cue onset. To investigate whether sleep influences the spatial distribution of these alpha effects, we compared the alpha-band time-frequency representations for successfuly recalled trials between pre-sleep and post-sleep recall phases during this.7 - 1.5 s time window. Results revealed two distinct spatial clusters (*p* <.05, two-sided, cluster-corrected) of alpha power differences between pre-and post-sleep recall (**Figure 4a**). Stronger pre-sleep alpha decreases were predominantly localised in left central-parietal regions, while stronger post-sleep alpha decreases shifted toward right anterior temporal regions. Importantly, no statistically significant differences between pre-and post-sleep recall emerged when contrasting unsuccessful recall trials, mitigating concerns about mere circadian effects underlying these results.

To further examine this shift in alpha decrease, we next tested whether N3 sleep predicts the observed shifts in recall networks. Using the previously defined (temporal, parietal) clusters from the pre-vs. post-sleep comparison (**Figure 4a**), we correlated the proportion of N3 sleep with the *change* in alpha decreases between pre-sleep and post-sleep (PM minus AM;.7-1.5 s). Here, a positive correlation would mean that more time spent in N3 sleep is associated with higher post-sleep alpha decreases. This analysis, shown in **Figure 4b**, revealed four significant relationships. Change in alpha decrease in the right temporal cluster was positively associated with i) time spent in N3 sleep (*r* =.47, *p* <.05) and ii) memory retention (*r* =.51, *p* <.05), aligning with the idea that sleep supports a functional shift in recall networks towards anterior temporal regions and that this shift is associated with higher levels of memory retention. The change in alpha decrease in the parietal cluster, on the other hand, showed a negative relationship with i) time spent in N3 sleep (*r* = -.49, *p* <.05) and memory retention (*r* =-.42, *p* <.05). No such relationships were found between any other sleep stage. This overall suggests that N3 sleep plays a role in restructuring memory retrieval processes, reinforcing temporal lobe involvement in post-sleep recall.

As mentioned in the introduction, a key prediction of systems consolidation models is that sleep fosters large-scale reorganisation of brain networks. To test this notion, we first derived memory networks by assessing alpha-band *i*-coherence during a 0.7–1.7s window for successful recall trials, separately for pre-sleep and post-sleep recall. We then quantified the global reorganisation of these networks using the cosine distance between the resulting connectivity matrices across the previously defined regions of interest (‘connectome distance’, see **Figure 4c** upper panels for a visual representation). Crucially, this analysis revealed a significant positive correlation between time spent in N3 sleep and the connectome distance between pre-sleep and post-sleep recall (**Figure 4c**, lower panel; *r* =.44, *p* =.047). This correlation also held when looking at the absolute duration of N3 sleep, *r* =.47, *p* =.026), indicating that greater time in N3 sleep was associated with stronger network reorganisation. Moreover, greater reorganisation of the alpha-band connectome was itself correlated with higher memory retention following sleep (*r* =.57, *p* =.003), suggesting that *behaviourally-relevant* overnight memory stabilisation is accompanied by large-scale changes in network connectivity. No such relationship was found with any other sleep stage, reinforcing the specificity of slow wave sleep in correlating with memory network reorganisation. Likewise, no comparable effects were seen for unsuccessful recall trials (all correlation *ps* >.25), again assuring that the results do not reflect time-of-day differences.

## Discussion

This study investigated the role of sleep in the transformation of episodic memory networks. Using EEG scalp and source analysis of an episodic memory task (**Figure 1**), we first replicated the finding of alpha power decreases for successful compared to unsuccessful recall between 0.7 to 1.5 seconds in the core recollection network (**Figure 2**) (Griffiths et al., 2019; Martín-Buro et al., 2020; Rugg & Vilberg, 2013; Staresina & Wimber, 2019; Treder et al., 2021). Turning to the impact of sleep on episodic memory, we observed the expected decline in recall performance overnight, albeit to different extents across participants (**Figure 3**). Importantly, contrasting post-sleep vs. pre-sleep recall networks, we observed a shift of alpha recall effects from central-parietal regions to the anterior temporal lobe (ATL; **Figure 4**). Time spent in N3 sleep not only predicted levels of memory retention but also correlated with the extent of this parietal-to-ATL shift across participants. Moreover, participants with better overnight memory retention displayed a more pronounced parietal-to-ATL shift. Lastly, examining brain-wide connectivity profiles, we found that time spent in N3 sleep predicted the extent of overnight connectivity changes during successful recall. This change in connectivity profiles in turn tracked levels of memory retention. Together, our findings thus unveil how slow wave (N3) sleep reshapes regional and brain-wide dynamics supporting episodic memory retrieval.

**Figure 3.**
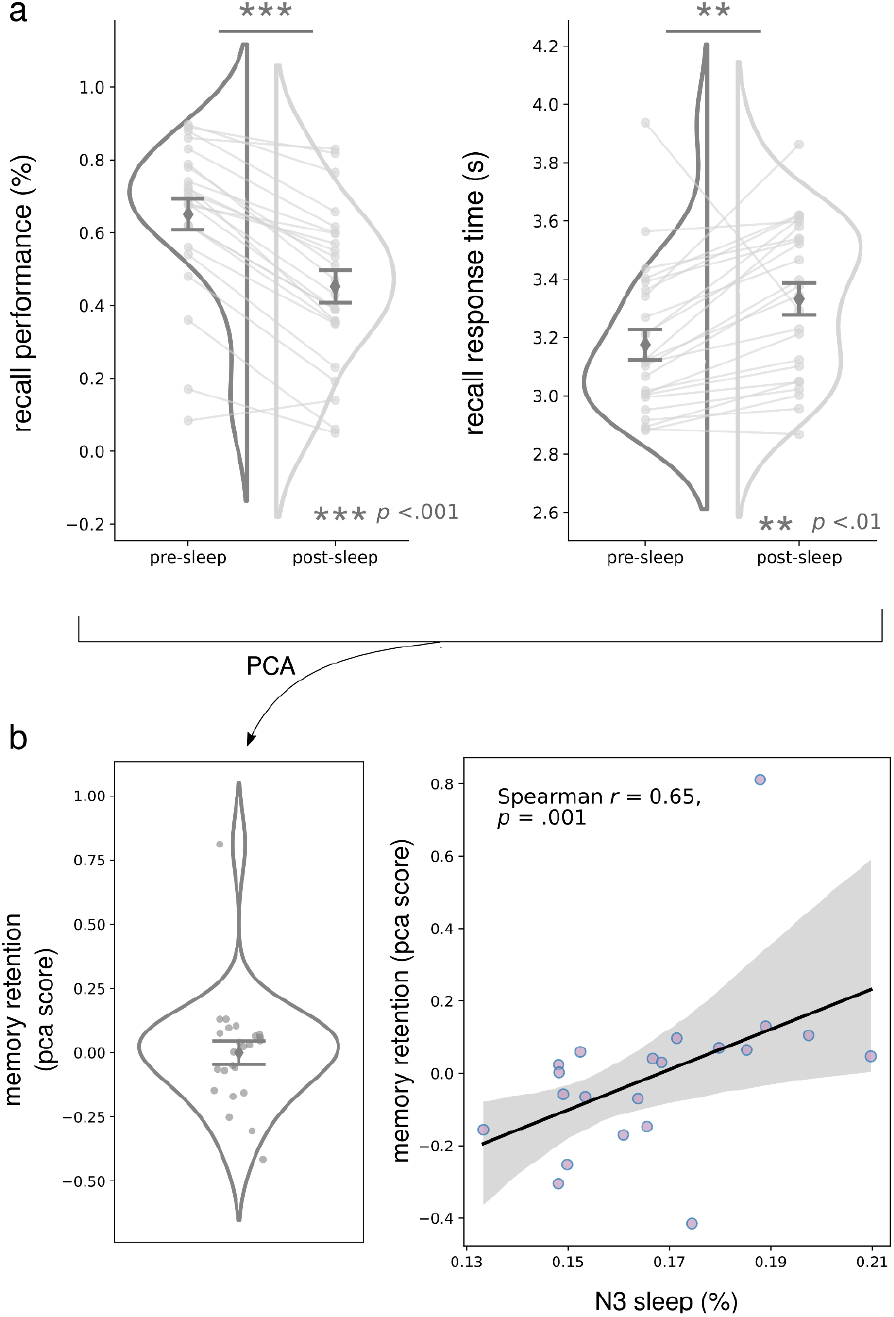
Recall performance and relationship with sleep. (a) Behavioural results showing changes in recall performance before and after sleep. The left panel displays adjusted recall accuracy (%), and the right panel shows recall response time (s) following the response prompt (which appeared 2.5 seconds after cue onset). Violin plots illustrate the distribution of individual data points, with mean and standard error bars overlaid. Each line links a participant’s pre - and post-sleep performance. A significant decrease in recall accuracy (*p* <.001) and an increase in recall response time (*p* <.01) were observed post-sleep, indicating overnight forgetting. (b) Principal Component Analysis (PCA) was applied to both behavioural measures (accuracy and response time) to derive a single memory retention score. The left panel shows the distribution of individual PCA scores, summarizing overnight memory retention. The right panel depicts a significant positive correlation between N3 sleep proportion and memory retention.

**Figure 4.**
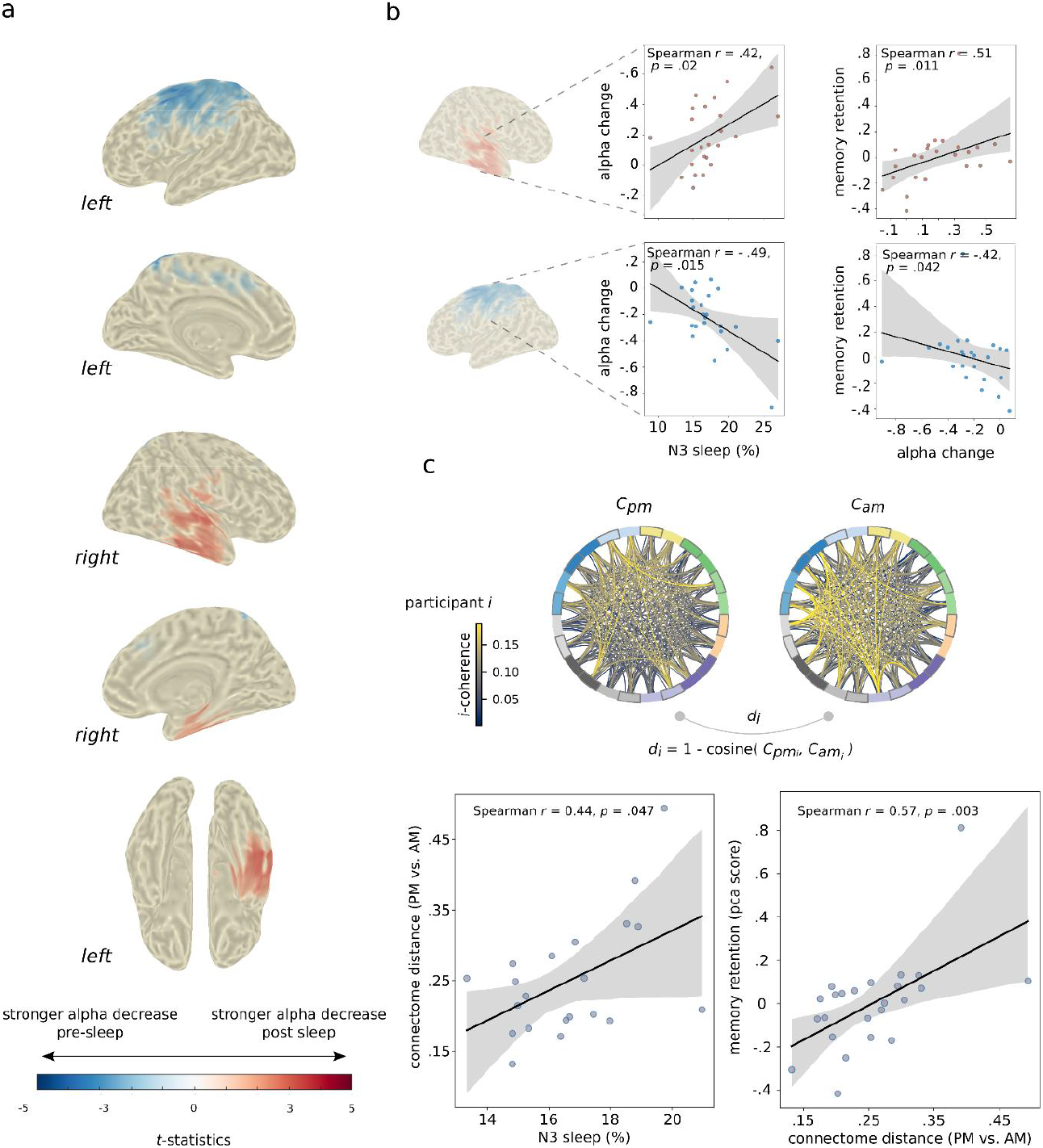
Pre-sleep vs. post-sleep alpha recall effects, and relationship with time-spent in N3 sleep. (a) Statistical comparison of alpha-band decreases between pre-sleep and post-sleep recall success. Blue clusters indicate regions where alpha decreases were stronger pre-sleep, while red clusters indicate stronger post-sleep alpha decreases (cluster-test, *p* <.05, two-sided;.7 - 1.5 s). (b) Correlation between the change in pre-vs. post-sleep alpha decreases for successful recall and time spent in N3 sleep was performed across participantsfor thetwo predefined region of interest shown in (a). Correlations shown in the first row indicatethat N3 sleep proportion wasassociated with stronger temporal lobeengagement (alpha power decreases) post sleep, which was in turn positively associated with memory retention. Correlations shown in the second row indicate that N3 sleep proportion was associated with less post-sleep parietal engagement, which was in turn negatively associated memory retention. (c) Network-level reorganisation of alpha-band connectivity acrosspre-and post-sleep recall. Circularplots illustrate alpha *i*-coherence connectivity patterns before (C_pm_) and after (C_am_) sleep for an example participant. Below, scatter plots show that greater N3 sleep proportion was associated with higher global connectome distance between pre-and post-sleep recall and that greaterconnectomereorganisation was in turn associated with better memory retention.

Models of systems consolidation propose a gradual shift from hippocampal to neocortical involvement during recall with time (Born & Wilhelm, 2012; Frankland & Bontempi, 2005; Klinzing et al., 2019; McClelland et al., 1995; Rasch & Born, 2013). Previous fMRI studies in humans have provided evidence for hippocampal-to-cortical shifts from immediate to delayed memory recall (Gais et al., 2007; Takashima et al., 2009), although the question whether hippocampal involvement in recall ever subsides remains debated (Barry & Maguire, 2019; Moscovitch et al., 2005; Nadel et al., 2000, 2007; Squire et al., 2015; Winocur et al., 2010). Nevertheless, converging evidence supports the notion that sleep (compared to wake) contributes to the reorganisation of retrieval networks. For instance, one study (Payne & Kensinger, 2011) examined emotional memory retrieval in the morning after 12 h of sleep compared to retrieval in the evening after 12 h of being awake. Sleep resulted in a shift from engagement of a diffuse memory retrieval network—characterised by widespread activity in lateral prefrontal and parietal cortices—to a more selective set of regions, including the amygdala and ventromedial prefrontal cortex. Effective connectivity analyses indicated enhanced connectivity among limbic regions following sleep compared to wakefulness. Sleep-related changes in connectivity profiles were also found in an fMRI study investigating motor sequence learning (Debas et al., 2014). Post-learning sleep (compared to post-learning wake) led to greater cortico-striatal connectivity after sleep. While N3/slow wave sleep in particular is considered to promote changes in retrieval networks (Diekelmann & Born, 2010; Gais & Born, 2004; Stickgold & Walker, 2005; Winocur & Moscovitch, 2011), these fMRI studies have not demonstrated a link between sleep architecture and the observed changes in recall activity. Conducting PSG between immediate and delayed recall, our data provide evidence for a direct link between N3 sleep and changes in local as well as brain-wide retrieval networks. Given the presence of both slow oscillations/slow waves and spindles (as well as hippocampal ripples) during N3, future studies should investigate whether these sleep rhythms and/or their coupling (Staresina et al., 2015; Staresina, 2024) confer the beneficial effect of N3 sleep on memory consolidation and transformation of recall networks. Indeed, recent work (Nicolas et al., 2025) using closed-loop targeted memory reactivation (TMR) has shown that by presenting auditory reminders during slow-oscillation UP states (and thereby eliciting stronger coupled spindle activity), connectivity increased within the task-relevant striato-motor networks from pre-to post-sleep.

As illustrated in **Figure 4a**, we observed a shift in alpha recall effects from pre-to post-sleep retrieval, migrating from left centro-parietal cortex to the right anterior temporal lobe (ATL). The involvement of parietal cortex during pre-sleep recall is consistent with previous findings on immediate memory retrieval (Vilberg & Rugg, 2008; Wagner et al., 2005) and may reflect enhanced fidelity of successfully recalled memories in immediate compared to delayed recall. Notably, our finding of increased ATL engagement post-sleep aligns with the results of Nieuwenhuis et al. (2012), who used MEG and connectivity analyses to demonstrate that delayed memory recall (25 h post-encoding) was associated with greater ATL recruitment. In agreement with Nieuwenhuis et al. (2012), we propose that this shift to the ATL might reflect a transformation of the neural representation of memory, perhaps reflecting increasing reliance on higher-level semantic features during delayed recall. This interpretation is consistent with prior work demonstrating the ATL’s critical role in semantic processing and conceptual integration (Patterson et al., 2007; Ralph et al., 2010; Ralph et al., 2017). Future research employing more advanced analytical approaches, such as encoding models (Nishimoto et al., 2011; Kay et al., 2008; Naselaris et al., 2011) and representational similarity analysis (Kriegeskorte et al., 2008; Cichy et al., 2014; Ritchie et al., 2019), will be essential for disentangling the specific computations underlying putative memory transformations.

## Data and code availability

Data and code for this project will be available publicly upon acceptance of the manuscript on OSF. DOI 10.17605/OSF.IO/GURCE

## Acknowledgements

This project has received funding from the European Research Council (ERC) under the European Union’s Horizon 2020 (grant agreement no. 101001121) awarded to B.P.S., as well as funding support from The Royal Society (project NIF\R1\221006) awarded to S.F.S.

## CRediT author contributions

Conceptualisation, B.P.S.; methodology, B.P.S.; software, B.P.S; project administration, P.P., S.F.S.; data curation, P.P and S.F.S.; validation, S.F.S.; formal analysis, S.F.S..; investigation, P.P. and S.F.S.; writing – original draft, S.F.S.; writing – review & editing, S.F.S., P.P., and B.P.S.; visualisation, S.F.S.; supervision, B.P.S.; funding acquisition, B.P.S., S.F.S.

## Supplementary material

**Supplementary Figure 1.**
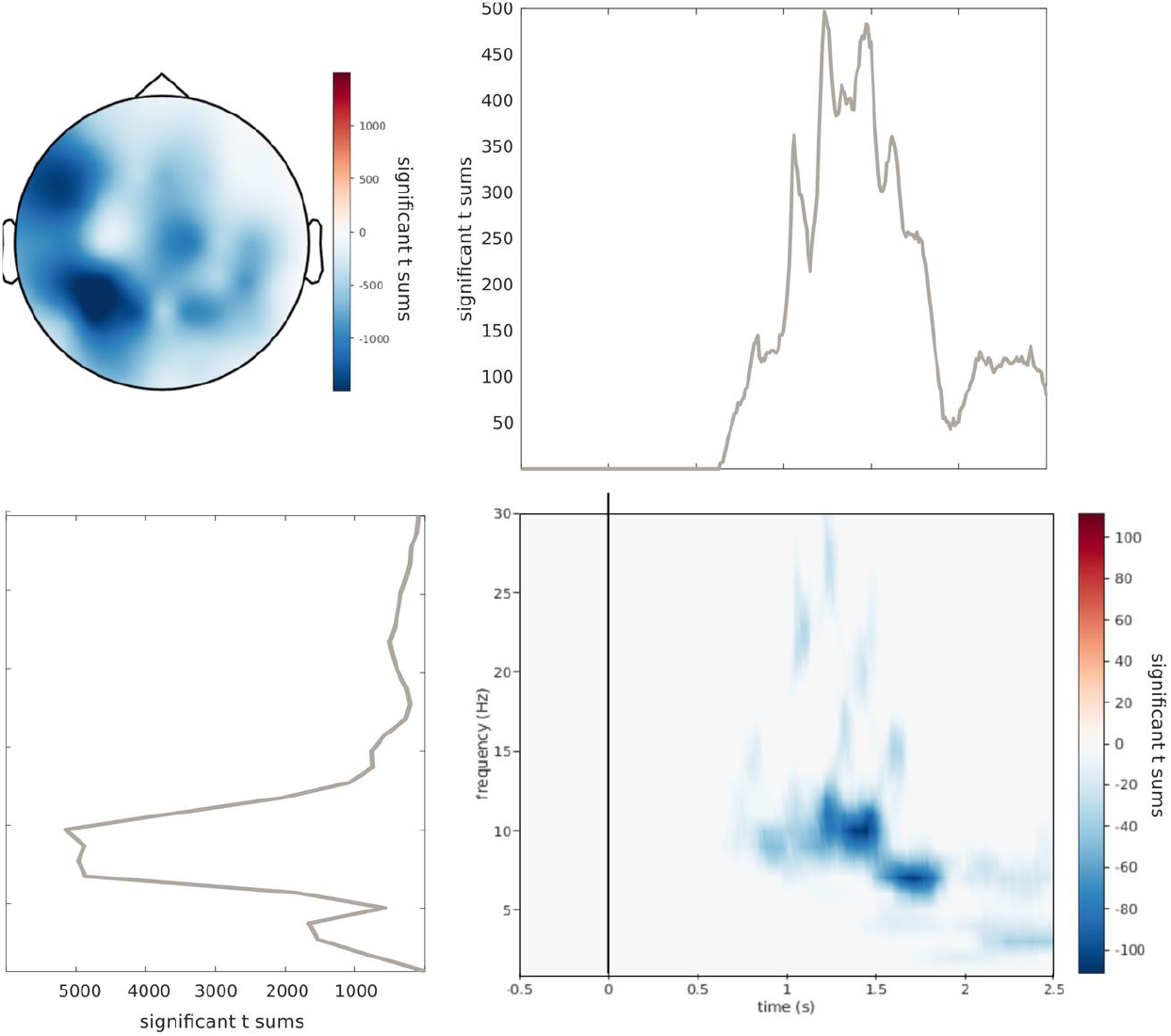
Cluster-based permutation test on 3D (time-frequency-channel) EEG data. We compared successful vs. unsuccessful recall conditions using a dependent samples t-test with cluster correction. The top-left panel shows the topographic distribution of significant t-sums, indicating spatial clusters of statistical differences. The top-right panel illustrates the time series of significant t-sums across trials, highlighting the most significant time windows. The bottom-left panel presents the frequency-wise distribution of significant clusters, illustrating spectral differences, peaking at 10Hz. The bottom-right panel displays the time-frequency representation of significant clusters, showing when and at which frequency effects emerge. Statistical parameters include 1000 random permutations, an alpha threshold of 0.05, and cluster correction using max-sum statistics (*p* =.008).

